# A photoswitchable positive allosteric modulator to control the activation of the metabotropic glutamate receptor 5 by light

**DOI:** 10.1101/2024.12.24.629646

**Authors:** Anaëlle Dumazer, Roser Borras-Tuduri, Iona Truong, Xiaojing Cong, Fanny Malhaire, Xavier Rovira, Xavier Gomez-Santacana, Laurent Givalois, Guillaume Lebon, Amadeu Llebaria, Cyril Goudet

**Affiliations:** IGF, Université de Montpellier, CNRS, INSERM, 34094 Montpellier, France; MCS, Laboratory of Medicinal Chemistry and Synthesis, Institute of Advanced Chemistry of Catalonia (IQAC-CSIC), Barcelona, Spain; MMDN, Univ Montpellier, EPHE-PSL, INSERM, Montpellier, France; Laval University, Faculty of Medicine, Department of Psychiatry and Neurosciences, CR-CHUQ, Québec city (QC), Canada

**Keywords:** Glutamate, GPCR, mGlu_5_, photopharmacology, photochromic ligand, photoswitch

## Abstract

The metabotropic glutamate receptor 5 (mGlu_5_) is widely expressed in the brain, where it plays an important role in synaptic plasticity, learning and memory, making it a therapeutic target of interest in various neurological disorders. In this study, we developed a photoswitchable positive allosteric modulator (PAM) of the mGlu_5_, as a novel tool for this clinically relevant drug target. To that aim, we used an azologisation strategy of the mGlu_5_ PAM agonist VU0424465 leading to the molecule azoglurax. We observed a reversible photoisomerization of azoglurax in solution with optimal wavelengths of 365 nm and 435 nm for *trans* to *cis* and *cis* to *trans* isomerization, respectively. In cell-based assays, azoglurax potentiates the agonist-induced activity of mGlu_5_ with a sub-micromolar potency in the dark. This potency is reduced under UV illumination. Similar to its parent molecule, azoglurax acts as an allosteric agonist of mGlu_5_, activating the receptor in absence of glutamate, as demonstrated on a glutamate-insensitive mutant receptor. Docking and site-directed mutagenesis experiments also suggest that azoglurax and VU0424465 bind the same pocket. In addition, molecular dynamics on *cis*-azoglurax-bound mGlu_5_ suggests that it azoglurax cis isomer does not bind stably in the receptor, in contrast to the *trans*-isomer, explaining the difference of activity between the two isomers. In conclusion, azoglurax is the first mGlu_5_ photoswitchable PAM agonist reported to date, retaining the properties and the binding mode of its parent in the dark, while the insertion of an azobenzene confers light-regulated activity.

**Graphical abstract.**
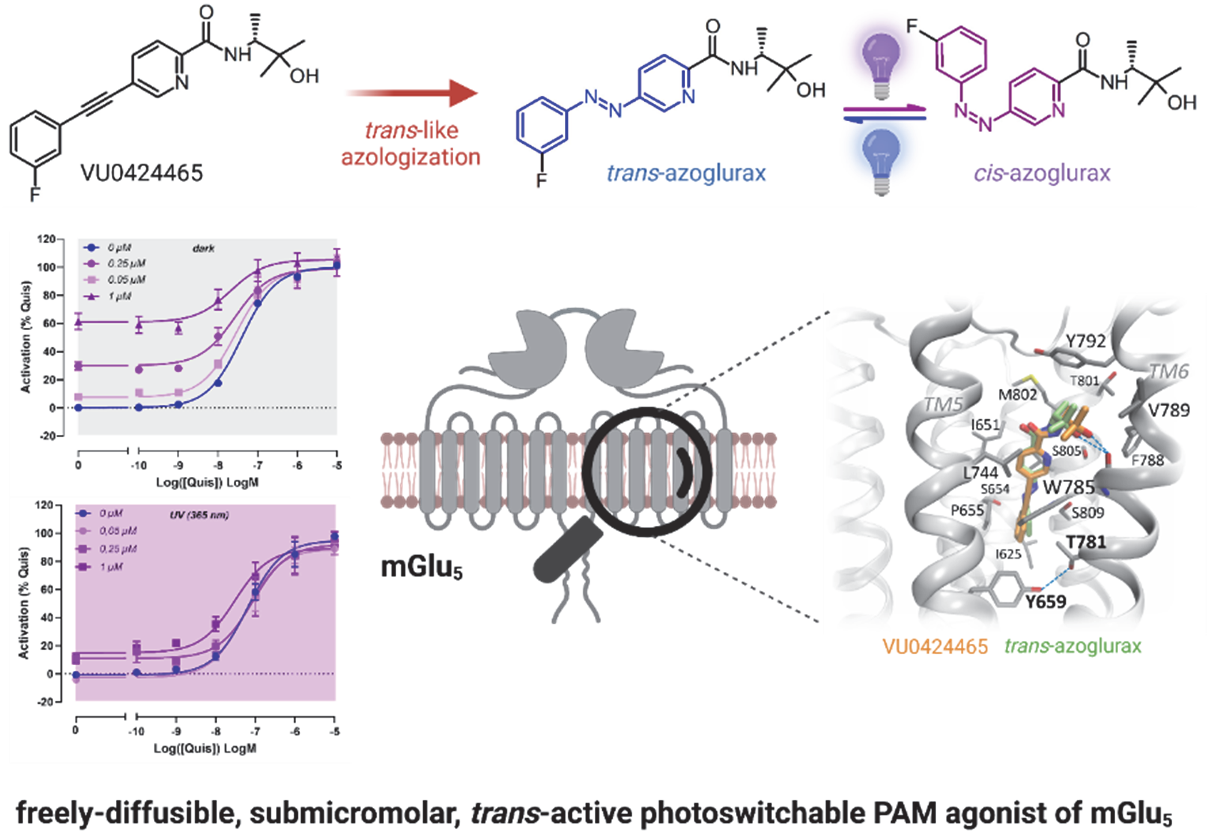

## 2. Introduction

Glutamate is the main excitatory neurotransmitter in the nervous system of adult mammals [1]. It carries out its functions by binding to two classes of receptors: ionotropic glutamate receptors [2], which are ligand-gated ion channels responsible for fast synaptic responses, and metabotropic glutamate receptors (mGluRs) [3], which are G protein-coupled receptors (GPCRs) responsible for the neuromodulatory actions of glutamate. The various mGluRs are widely expressed in the nervous system, where they modulate glutamatergic transmission and thus contribute to many important physiological processes [4]. Many neurological disorders are caused by abnormalities in glutamatergic transmission, making mGluRs targets of therapeutic interest [5–7].

Eight different mGluRs subtypes have been identified, named mGlu1 to mGlu8, subdivided into three groups based on sequence homology, signaling and pharmacology [3]. Group I consists of mGlu_1_ and mGlu_5_, group II of mGlu_2_ and mGlu_3_, while group III comprises mGlu_4_, mGlu_6_, mGlu_7_ and mGlu_8_. The structure of mGluRs consists of a large extracellular *N*-terminal domain where glutamate binds, called the Venus flytrap (VFT) domain, which is linked by a cysteine-rich domain (CRD) to the 7-transmembrane alpha-helical domain (7TM) typical of all GPCRs [8–12]. mGluRs are obligate dimers, with the two protomers cross-linked by a disulfide bridge located at the top of the extracellular domain. They can form homodimers and heterodimers [13]. Glutamate binding in the extracellular domain induces a cascade of molecular events within the dimer, which ultimately leads to the coupling of the active 7TM domains to the G protein and the initiation of the cellular response [14–16].

As the L-glutamate binding site is highly conserved between different mGluRs, it is difficult to identify subtype-selective orthosteric ligands [7]. To circumvent this problem, most recent efforts have focused on the development of allosteric modulators, which bind to topographically distinct sites from the orthosteric site. Most allosteric sites are located in the 7TM region of mGluRs, which show greater sequence divergence between subtypes and therefore offer greater selectivity. Allosteric modulators can increase or decrease the affinity or potency of orthosteric ligands and are referred to as positive or negative allosteric modulators (PAM or NAM). In addition, PAMs that have intrinsic efficacy are referred to as PAM agonists (abbreviated ago-PAM herein) [7, 17, 18].

Among the mGluRs family, the mGlu_5_ subtype is an important modulator of synaptic plasticity involved in cognitive functions. It is widely distributed in the brain, notably in cortex, hippocampus, basal ganglia and thalamus [4, 19]. In neurons, mGlu_5_ receptor is predominantly found in postsynaptic elements within the central nervous system, where it increases neuronal excitability and membrane depolarization following activation by glutamate. It is also expressed on glia, including astrocytes and microglia, where its activation is important to glia function and interaction with nerve cells [20]. mGlu_5_ preferentially couples to the G_q/11_ proteins, mediating signalization through diacylglycerol, inositol 1,4,5-trisphosphate and intracellular Ca^2+^. Given the high therapeutic value of mGlu_5_, this receptor has been the focus of intense development of allosteric modulators, resulting in the existence of a large number of NAMs and PAMs [3]. Preclinical studies have highlighted the therapeutic potential of mGlu_5_ blockade by NAMs on neurological and neurodegenerative diseases, such as anxiety [21], depression [22], chronic pain [23], Parkinson and Alzheimer diseases [24]. On the other hand, there is great potential to alleviate various symptoms of schizophrenia through mGlu_5_ PAMs.

Photopharmacology is an emerging strategy based on the use of small, diffusible ligands whose interaction with their target is controlled by light and therefore does not require genetic manipulation, unlike optogenetics [25]. It is a powerful fundamental research tool which, combined with optical technologies, enables the investigation of endogenous biological processes with micrometric spatial resolution and millisecond temporal resolution. This high spatiotemporal resolution is particularly interesting for targets such as mGlu_5_, which are widely expressed [4]. Two major categories of freely-diffusible photosensitive ligands are available: photoactivable ligands (also named caged ligands)[26], which are irreversibly activated by light, and photoswitchable ligands [27], which can be reversibly switched from an active to an inactive conformation. Photopharmacology has been applied to many GPCRs [28–30], including mGlu_5_. The first photoswitchable allosteric modulator of GPCR was Alloswitch-1, a NAM of mGlu_5_ [31]. It was subsequently derived into a series of analogues that enabled the structure-activity relationships of this family of compounds to be explored [32, 33]. Finally, there is also a caged NAM, JF-NP-26, derived from raseglurant [34]. However, to date, there is no photosensitive ligand that can activate or potentiate this receptor.

Here we aimed to develop a photoswitchable mGlu_5_-positive allosteric modulator as a novel tool for this clinically relevant drug target. Recent cryoEM structure of mGlu_5_ bound to the high affinity ago-PAM VU0424465 revealed a binding mode very similar to the NAM M-MPEP and to the photoswitchable NAM alloswitch-1, the VU0424465 ethynyl group’s aligning very well with alloswitch-1 azo group M-MPEP ethynyl [12, 35]. From these observations, we identified VU0424465 as a candidate for developing a first photoswitchable mGlu_5_ PAM. The VU0424465 [36] has been modified by an azologization process and the resulting ligand (named azoglurax) photochemical and photopharmacological properties were characterized. Its binding mode in *trans* and possible *cis*- configuration were further investigated by using combining photopharmacology and alanine mutant localized in the 7TM of the mGlu_5_ receptor and molecular dynamic simulation. Taken together, our results demonstrate that azoglurax is the first photoswitchable PAM of the mGlu_5_ receptor, retaining the properties of ago-PAM and the binding mode of its parent in the dark, while the insertion of an azobenzene confers light-regulated activity.

## 3. Materials and methods

### 3.1. Synthetic methods and synthesis procedures

All chemicals and solvents were obtained from commercial suppliers and used without purification. DMF was dried by passing through a PureSolv solvent purification system. Compounds **2** and **5** were synthesised according the procedures detailed below. Reactions were monitored by thin layer chromatography (Macherey-nagel ALUGRAM SIL G/UV_254_) by visualisation under 254 nm lamp. Flash column chromatography was performed manually with Silica Gel 60 (40-63 µm, PanReac) or automated with SNAP KP-C18-HS (12g, 50 µm, Biotage) cartridge on Isolera One with UV-Vis detection (Biotage). Nuclear magnetic resonance (NMR) spectra were determined with and on a 400 MHz Brüker Avance NEO instrument. Chemical shifts are reported in parts per million (ppm) against the reference compound using the signal of the residual non-deuterated solvent (CDCl_3_ δ = 7.26 ppm (^1^H), δ = 77.16 ppm (^13^C); CD_3_OD δ = 2.50 ppm (^1^H), δ = 39.52 ppm). NMR spectra were processed using MestreNova 10.0.2 software (Mestrelab Research). The peak multiplicities are defined as follows: s, singlet; d, doublet; t, triplet; q, quartet; dd, doublet of doublets; ddd, doublet of doublets of doublets; dt, doublet of triplets; dq, doublet of quartets; td, triplet of doublets; tt, triplet of triplets; tdd triple of doublets of doublets; br, broad signal; m, multiplet; app, apparent. Purity determination was performed with Liquid Chromatography (LC) coupled to a photodiode detector (PDA) and a mass spectrometer (MS) with integration at 230 nm using a Waters 2795 Alliance separation module coupled to a diode array detector (Agilent 1100) scanning at a wavelength range of 210-600 nm and an ESI Quattro Micro MS detector (Waters) in positive mode with mass range (m/z) of 150-1500. A column ZORBAX Extend-C18 3.5 μm 2.1×50mm (Agilent) at 35°C was used with a mixture of A = H_2_O + 0.05% formic acid and B = MeCN + 0.05% formic acid as mobile phase and the method as follows: flow 0.5 mL/min, Gradient t = 0.0 min 5% B, t = 0.5 min 5% B, t = 5.5 min 100% B, t = 7.0 min 100% B, t = 8.0 min 5% B, t = 10.0 min 5% B, total runtime: 10 min or. High resolution mass spectra (HRMS) and elemental composition were analysed by FIA (flux injected analysis) with Ultrahigh-Performance Liquid Chromatography (UPLC) Aquity (Waters) coupled to LCT Premier Orthogonal Accelerated Time of Flight Mass Spectrometer (TOF) (Waters). Data from mass spectra were analysed by electrospray ionisation in positive and negative mode. Spectra were scanned between 100 and 1500 Da with values every 0.2 seconds and peaks are given m/z (% of basis peak). Data was acquired with MassLynx software version 4.1 (Waters) and analyses were performed at the mass spectroscopy service of IQAC-CSIC.

#### *(E*)-5-((3-fluorophenyl)diazenyl)picolinic acid (5)

A solution of 5-aminopicolonic acid (**3**, 100 mg, 0.72 mmol) in toluene (0.7 mL) and 1-fluoro-3-nitrosobenzene (**4**, 136 mg, 1.09 mmol) and acetic acid (4.0 mL) was added. The resulting mixture was stirred at 50 °C for 4 days. The solvent was removed in vacuo and the resilting residue was purified by automated reverse phase flash chromatography on C18 (H_2_O/MeCN + 0.05% formic acid 2:3 to 0:1) to give the title product **5** as an orange solid (66 mg, 37% yield). LC-MS: *cis*-**5** RT = 3.00 min, λ_max_ = 280, 420 nm, [M+H]^+^ (m/z) = 246.32, purity 3% (230 nm); *trans*-**5** RT = 3.44 min, λ_max_ = 324 nm, [M+H]^+^ (m/z) = 246.25, purity 90% (230 nm); ^1^H NMR (400 MHz, CD_3_OD) δ 9.19 (app. s, 1H), 8.40 (dd, *J* = 8.4, 2.0 Hz, 1H), 8.34 (d, *J* = 8.4 Hz, 1H), 7.89 (dt, *J* = 8.0, 1.3 Hz, 1H), 7.70 (dt, *J* = 9.7, 2.2 Hz, 1H), 7.64 (td, *J* = 8.1, 5.8 Hz, 1H), 7.36 (tdd, *J* = 8.3, 2.6, 0.9 Hz, 1H). ^13^C NMR (101 MHz, MeOD) δ 167.0, 165.9, 163.5, 155.3, 155.3, 151.1, 150.6, 146.9, 132.2, 132.1, 130.4, 127.1, 122.4, 120.4, 120.2, 109.1, 108.9. Correlations performed by COSY, HSQC, HMBC (see supporting information).

#### (*R*,*E*)-5-((3-fluorophenyl)diazenyl)-N-(3-hydroxy-3-methylbutan-2-yl)picolinamide (2)

Triethylamine (0.37 mL, 2.7 mmol) was added to a solution of the carboxylic acid **5** (66 mg, 0.27mmol), the salt **6** (94 mg, 0.43 mmol) and HATU (307 mg, 0.81 mmol) in dry DMF (2.7 mL) and the resulting solution was stirred at rt for 18 h. The solution was diluted in AcOEt (50 mL), washed with water (25 mL) and sat. NaCl (25 mL), dried over Na_2_SO_4_ and the solution was concentrated in vacuo. The resulting residue was purified by normal phase flash chromatography on silica gel (DCM/EtOAc 1:1) to give the title product **2** as an orange solid (40 mg, 45% yield). LC-MS: *trans*-**2** RT = 3.88 min, λ_max_ = 326 nm, [M+Na]^+^ (m/z) = 353.25 [M+H]^+^ (m/z) = 331.33, [M-OH]^+^ (m/z) = 313.33, purity 99% (230 nm). HRMS [M+H]^+^ (m/z) calculated for C_17_H_20_FN_4_O_2_ ^+^ 331.1565, found 331.1569. ^1^H NMR (400 MHz, CDCl_3_) δ 9.10 (dd, J = 2.2, 0.7 Hz, 1H), 8.35 (dd, J = 8.4, 0.8 Hz, 1H), 8.29 (br, 1H), 8.26 (dd, J = 8.4, 2.3 Hz, 1H), 7.81 (ddd, J = 7.9, 1.8, 1.0 Hz, 1H), 7.64 (ddd, J = 9.6, 2.6, 1.7 Hz, 1H), 7.53 (td, J = 8.1, 5.8 Hz, 1H), 7.24 (tdd, J = 8.2, 2.5, 1.1 Hz, 1H), 4.16 (dq, J = 9.2, 6.9 Hz, 1H), 1.32 (d, J = 6.9 Hz, 3H), 1.32 (s, 3H), 1.30 (s, 3H). ^19^F NMR (376 MHz, CDCl_3_) δ −111.5.^13^C NMR (101 MHz, CDCl_3_) δ 164.6, 163.9, 162.2, 154.0, 153.9, 151.5, 149.0, 145.8, 130.7, 130.6, 128.5, 123.2, 121.4, 121.3, 119.2, 119.0, 108.5, 108.3, 73.3, 54.0, 27.8, 25.9, 16.1. Correlations performed by COSY, HSQC, HMBC (see supporting information).

### 3.2. Photochemistry

#### UV-Vis absorption spectroscopy

The UV-Vis absorption spectrum of samples of azoglurax (25µM, 20% DMSO in PBS) were measured between 270 and 550 nm with 2 nm fixed intervals in a quartz cuvette (105-200-85-40, Hellma) with an Evolution 350 UV-Vis spectrophotometer (Thermo Scientific). Illumination was achieved using LED illumination systems (CoolLED pE-4000), set at 100% power which corresponds to the following light-power for each wavelength: 365 nm: 100 mW, 385 nm: 160 mW, 405 nm: 100 mW, 435 nm: 25 mW, 460 nm: 125 mW, 470 nm: 35 mW, 500 nm: 25 mW. The LED was directed vertically to the cuvette at 1cm above the level of the liquid while protecting the cuvette from light. For the isomerization cycle experiment, the sample was prepared in pure DMSO at 25 µM with the 36 5nm-LED set at 50 mW and the 460 nm-LED set at 25 mW. The change of the wavelengths was monitored manually directly from the control board of the CoolLED system every 90 s.

#### PSS determination by LC/MS

The estimation of the isomer proportion at the photostationary state (PSS) after the corresponding illumination conditions was obtained with an 800 µL sample of azoglurax (100µM in PBS/DMSO 1:1). The sample was illuminated for 6 minutes with a 365 nm LED light (CoolLED pE4000) set at 100 mW and 200 µL were placed in a LC vial with insert for LC-MS analysis. To check that the PSS was reached after 6 minutes, it was illuminated for 4 additional minutes and analyzed again. Then, the same sample was illuminated with 460nm LED light (125mW) for 6 minutes and was analyzed by LC/MS. The percentage of *trans*- and *cis*-isomers was obtained from the integration of the two peaks corresponding to each isomer in the chromatogram displayed at 384 nm that corresponds to the wavelength of one *trans-cis* isosbestic point, where both isomers have the same molar extinction coefficient.

### 3.3. Cell culture & transfection

HEK293 cells (ATCC-CRL1573) were cultured in Dulbecco’s Modified Eagle Medium (DMEM, Life Technologies, Cergy Pontoise, France), supplemented with 10% of Fetal Bovine Serum (FBS), at 37°C and 5% CO_2_. Cells were regularly tested to be mycoplasma free. They were transiently transfected by electroporation with rat mGlu_5_ receptor wt or mutants. Receptors were also co-transfected with the glutamate transporter EAAC1 to minimize influence of extracellular glutamate. Cells were seeded in a poly-ornithine coated 96-well plate at the density of 100 000 cells/well. Plates were kept in the incubator at 37°C and 5% CO_2_ for 24 hours. The following day, the cell medium was replaced by Glutamax (Life Technologies, Cergy Pontoise, France) to reduce extracellular glutamate concentration and 2 hours before experiments.

### 3.4. Site-directed mutagenesis

mGlu_5_ mutants in the orthosteric binding site (mGlu_5_ Y236A+D318A, to be called mGlu_5_ YADA in the text) or in the allosteric binding site (mGlu_5_ Y659A; T781A; W785A; F788A; Y792A; S809A) were generated by site-directed mutagenesis and were verified by sequencing [9, 12]. All mutant receptors contain an *N*-terminal SNAP-tag to estimate their cell surface expression, by measuring fluorescence at 620 nm following covalent labeling of the receptor with SNAP-Lumi4-Tb. All mutant receptors were expressed at the cell surface and were functional. Transfection conditions were adjusted to yield similar cell surface expression than the WT mGlu_5_ receptor, as previously described [9, 12].

### 3.5. Cell-based pharmacological assays

The IP-One HTRF kit (Revvity) was used to estimate mGlu_5_ activity for the direct quantitative measurement of inositol monophosphate (IP_1_) in HEK293 cells transiently transfected with the human mGlu_5_ receptor, wild type or bearing the mutations described above. Prior to the experiment, the medium was changed to DMEM glutamax and cells were incubated for an additional 90 minutes to lower extracellular glutamate concentration. The experiments were carried out in 96-well plates, following the manufacturer’s conditions, adapted as described below. Cells were stimulated either with different concentrations of azoglurax or VU0424465 to induce IP accumulation in presence or absence of 10 nM of the agonist quisqualate (EC_20_) or with different concentrations of quisqualate in presence of fixed doses of azoglurax. Cells were incubated for 25 min, at 37 °C. After the stimulation, the medium was removed from the plate and 28µL of lysis buffer were added to each well. Cell lysates (14 µL) were transferred to a white 384-well plate in which d2-labeled IP analog and terbium cryptate-labeled anti-IP antibodies (6 µL) were added using a repetitive pipette and the plate was incubated for 1 hour. The fluorescence at 665 and at 620 was read with a pherastar HTRF HTS microplate reader (BMG Labtech). Afterwards, the HTRF ratios (665/620 nm) were transformed to IP concentration produced by the cells using a standard IP_1_ curve. For experiment in light conditions, cells were placed over a 365 nm-LED array (Teleopto) driven by a 96-LED Array Driver LAD-1 and monitored by a stimulator (STO mk II) set at 8 mW with a pulse frequency of 10 Hz.

### 3.6. Molecular modeling

Molecular modeling was based on the cryoEM structure of mGlu_5_ bound to VU0424465 [12]. Only the 7TM domain was considered to optimize the computational costs. *trans*-azoglurax was docked to the allosteric pocket using Autodock Vina [37], which returned a similar binding pose to VU0424465. *Cis*-azoglurax was obtained by switching the N=N double bond of *trans*-azoglurax in the docked model. The three ligand-7TM complexes were then subject to all-atom molecular dynamics (MD) simulations in an explicit membrane-solvent environment as follows.

#### MD system setup

CHARMM-GUI was used to assign the side-chain protonation states and embed the complex models in a bilayer of POPC lipids and cholesterol in a 3:1 ratio. The systems were solvated in a periodic box of explicit water and neutralized with 0.15 M of Na^+^ and Cl^-^ ions. We used the Amber ff14SB [38], GAFF2 [39] and lipid21 [40] force fields, the TIP3P water model and the Joung-Cheatham ion parameters [41]. Effective point charges of the ligands were obtained by RESP fitting of the electrostatic potentials calculated with the HF/6-31G* basis set using Gaussian 16 [42]. Additional force field parameters for *cis*-azoglurax were adopted from Böckmann *et al.* [43].

#### MD simulations

After energy minimization, MD simulations were carried out using Gromacs 2022 patched with Plumed 2.3. The LINCS algorithm was applied to constrain bonds involving hydrogen atoms, allowing for a time step of 2 fs. Each system was gradually heated to 310 K and pre-equilibrated during 10 ns in the *NPT-*ensemble. The REST2 technique [44] was then used to enhance the MD sampling in the *NVT* ensemble. REST2 is a type of Hamiltonian replica exchange simulation scheme that can capture GPCR conformational dynamics at the millisecond timescale with only microseconds of simulations (e.g. in our previous work [45–48]). REST2 performs many replicas of the same MD simulation system in parallel with the original system. The replicas have modified Hamiltonian to facilitate conformational changes. By frequently swapping the replicas and the original system during the MD, the simulations “travel” on different free energy surfaces and easily visit different conformational zones. Finally, only the samples in the original system (with unmodified free energies) are collected. The replicas are used only to help overcome the energy barriers. In REST2, the Hamiltonian of the replicas are modified by scaling (reducing) the force constants of the “solute” molecules in the simulation system. In this case, the protein and the ligands were considered as “solute”–the force constants of their van der Waals, electrostatic and dihedral terms were subject to scaling–in order to facilitate their conformational changes. The “effective temperatures” used here for generating the REST2 scaling factors ranged from 310 K to 750 K for 64 replicas, following a distribution calculated with the Patriksson-van der Spoel approach [49]. Exchange between the replicas was attempted every 1000 simulation steps. This setup resulted in an average exchange probability of ∼40 % throughout the simulation course. We performed 60 ns × 64 replicas (3.84 µs) of REST2 MD simulations for each system, discarding the first 10 ns for equilibration. The simulation trajectories on the original unmodified free energy surface were reassembled and analyzed.

### 3.7. Ligands

All chemicals were reagent grade (Merck, or Sigma, Germany or Fluorochem, Ireland). Quisqualate and Glutamate were purchased from Bio-techne Tocris. VU0424465 was synthetized as detailed in Nasrallah et al. [50]. Azoglurax (**2**) was synthesized as described in section 3.1.

### 3.8. Data analysis and statistics

Experimental data of azoglurax and quisqualate-mediated responses were fitted using the Hill equation and a simplified model of allosterism (eq. 1) including the minimum number of parameters [51] since the structure of the experimental data precludes the use of complete expressions of operational models. The usefulness of this procedure was evaluated in previous studies [52–54].

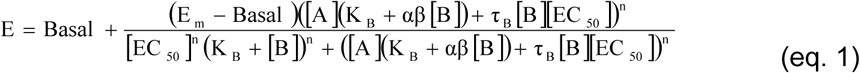

Where E_m_ is the maximum effect of the system; n, a parameter related with the slope of the curves; [A] and [B], the concentrations of the agonist A and the allosteric modulator B, respectively; τ_A_and τ_B_, the operational efficacies of A and B, respectively; K_A_ and K_B_, the dissociation constants of A and B, respectively; and α and β, the cooperativity parameters governing binding and function, respectively. Values of α and β greater, equal and lower than 1 indicate positive, neutral and negative cooperativity, respectively.

All data are reported as mean ± standard error of the mean (± SEM). Number of experiments and statistical tests that were performed on datasets are indicated in Figure Legends and/or in Tables. Data were analyzed with Microsoft excel 2016 and GraphPad Prism software v10, using one-way analysis of variance (ANOVA) and the appropriate post-hoc tests for multiple comparisons. Data were considered significant when p<0.05. P values less than 0.05 are indicated with one asterisk while P values less than 0.01, 0.001 and 0.0001 are indicated with two, three and four asterisks, respectively.

## 4. Results

### 4.1. Design and synthesis of the photoswitchable compound Azoglurax

A photoswitchable mGlu_5_ PAM has been designed following the azologization strategy of a diarylacetylene *trans*-like azostere motif. The replacement of the ethynyl group of, VU0424465 (**1**), a potent and selective mGlu_5_ ago-PAMs well described in the literature [36], with an azo bond led to the generation of the compound named azoglurax (**2**) **(Figure 1)**. The first step consisted in a Mills reaction between the commercial 5-aminopiconilic acid (**3**) and 1-fluoro-3-nitrobenzene (**4**) in acidic conditions to give the azo-carboxylic acid (**5**) in moderate yields. The activation of the carboxylic acid (**5**) with HATU and triethylamine and the following nucleophilic attack with the amine (**6**) gave the final product azoglurax (**2**) with moderate to good yields **(Figure 1)**.

**Figure 1:**
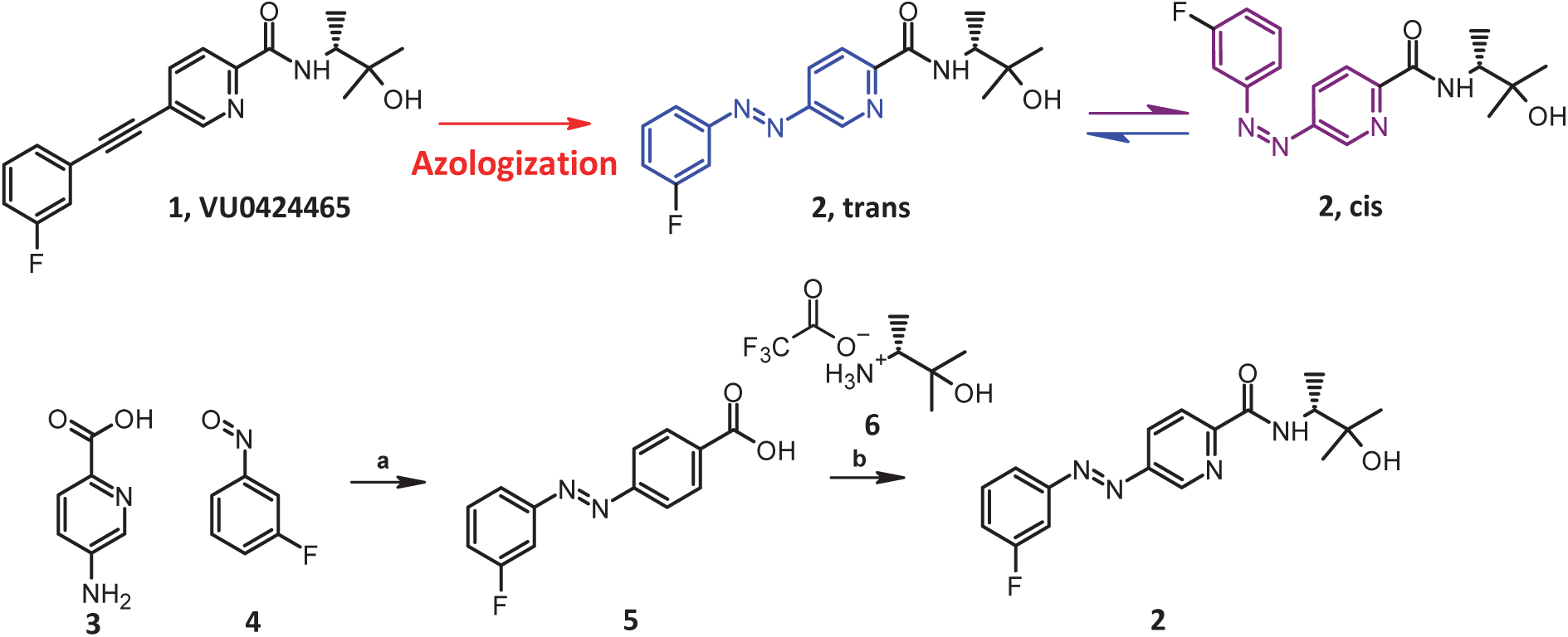
Design and synthesis of the photoswitchable mGlu_5_ PAMs azoglurax. **A.** Azoglurax was designed via the azologization of the well-known mGlu_5_ PAMs VU0424465. **B.** Synthetic route to obtain azoglurax. Reactants and conditions: (a) Toluene/AcOH, 50°C, 4 days, 37% (b) HATU, TEA, DMF, rt, 16h, 45-76%.

### 4.2. Reversible photoisomerization of azoglurax

We examined if, as expected, the inclusion of an azobenzene group within the chemical structure of VU0424465 confers a photochromic activity to azoglurax.

To determine its photochemical properties, we measured the UV−visible absorption spectrum of a solution of azoglurax at 25 µM in phosphate-buffered saline (PBS) with 20% dimethyl sulfoxide (DMSO), in the dark or after 3 minutes illumination at different wavelengths, ranging from 365 to 550 nm **(Figure 2A)**. The compound presents the typical azobenzene profile with high absorbance band around 320-335 nm corresponding to the π-π* transition of the *trans* isomer. After illumination with UV light (λ = 365 nm), we observed a decrease of the absorbance of this band, indicative of a conversion of the molecules to a photostationary state with a high proportion of *cis* isomer. On the other hand, illumination with blue-cyan light induced a photostationary state with a high proportion of *trans* isomer. The optimal wavelength for *trans* to *cis* isomerization is 365 nm while the optimal wavelength for *cis* to *trans* isomerization is 460 nm **(Table 1)**.

**Figure 2.**
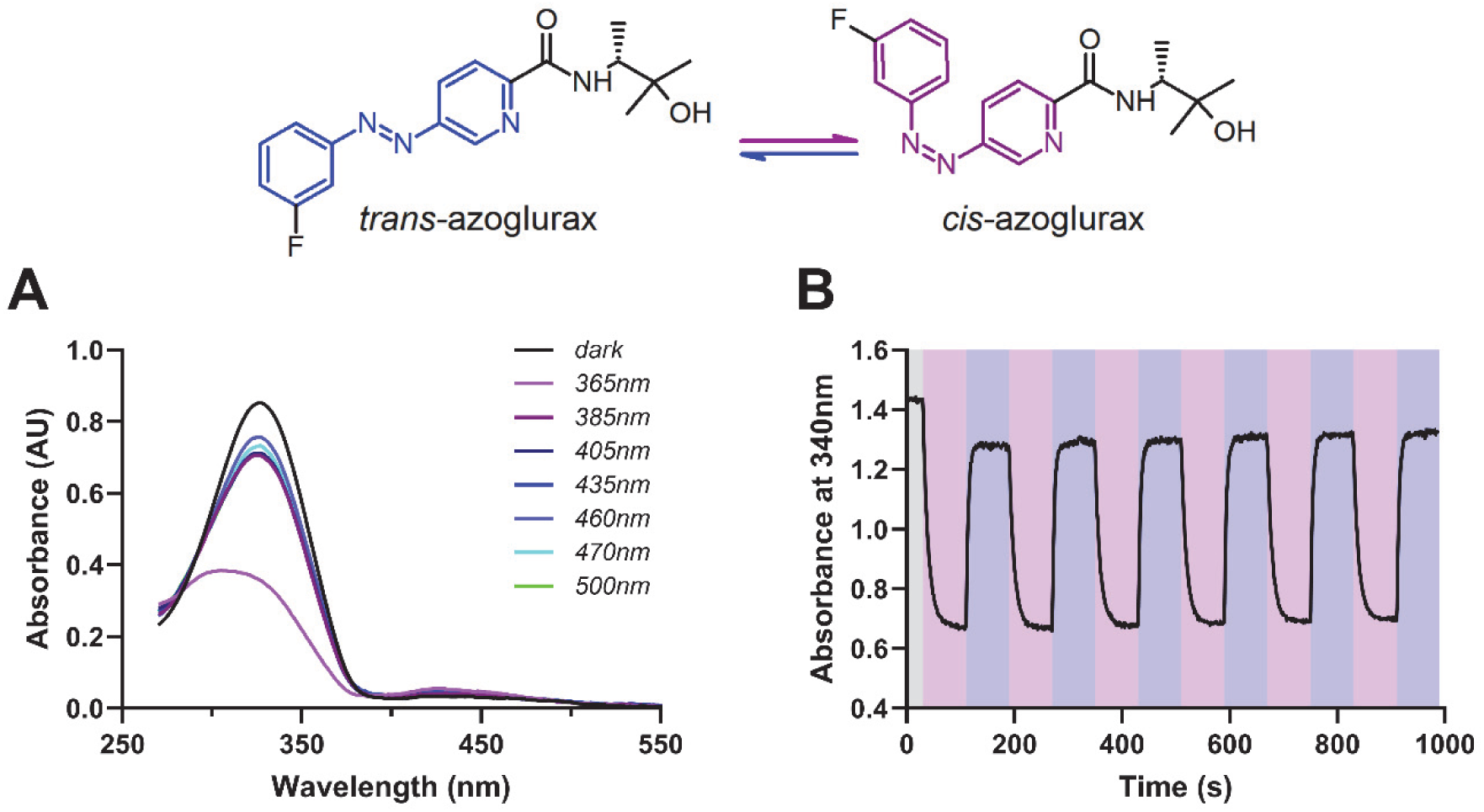
Photopharmacological characterization of azoglurax. Top. Scheme of *trans*/*cis* photoisomerization of azoglurax. **A.** UV/Vis absorption spectra of azoglurax (25 μM in PBS (20% DMSO) at 37°C under dark (black) and different light conditions for 3 minutes. **B.** UV/Vis absorption measurements of 25 μM azoglurax in DMSO at 340nm after repeated cycles of illumination with 365 nm (violet) and 460 nm light (blue).

**Table 1:** Photochemical properties of azoglurax. t_½_ *trans* -> PSS_365_ was determined using light power of 50 mW and t_½_ PSS_365_ -> PSS_460_ using a power of 25 mW using a 25µM sample in PBS (20% DMSO) at 37°C. The PSS_365_ has been determined with 100 µM samples in PBS/DMSO 1:1 and a 365 nm LED light of 100 mW and PSS_460_ with a 460nm LED light of 125 mW.

| $\lambda_{\max}$ (π-$\pi^*$) *trans* (nm) | $\lambda_{\max}$ ($n$-$\pi^*$) PSS<sub>460</sub> (nm) | PSS<sub>365</sub> (% *cis*) | PSS<sub>460</sub> (% *trans*) | $t_{1/2}$ *trans* → PSS<sub>365</sub> (s) | $t_{1/2}$ PSS<sub>365</sub> → PSS<sub>460</sub> (s) |
| --- | --- | --- | --- | --- | --- |
| 327 | 438 | 70 | 84 | 6.7 | 2.6 |

To determine the *cis/trans* isomeric proportion, the photostationary state of azoglurax was analyzed by HPLC after ten minutes of illumination (100 μM in PBS/DMSO 1:1) following UV-light irradiation (365 nm; trans to *cis* conversion) or blue light illumination (460 nm; *cis* to trans conversion). After 365 nm illumination, we observed a conversion from the *trans t*o the *cis*-isomer with a photostationary state (PSS_365_) containing 70% *cis*-isomer **(Table 1)**. Illumination at 460nm induces a reverse photoisomerization from *cis* to trans with a PSS_460_ containing 84% of the *trans* isomer **(Table 1)**.

The kinetic properties of light-induced photoisomerization were further characterized by measuring the absorption of a solution of azoglurax (25 µM in PBS 20% DMSO) at 30 nm after repeated cycles of 90 s illumination at 365 nm (50 mW) and 90 s at 460 nm (25 mW) **(Figure 2B)**. The *trans* to *cis* isomerization in the sample and illumination conditions described is relatively fast with a half-life to reach to the PSS_365_ of 6.7 s after illumination at 365 nm. The *cis* to *trans* isomerization is also fast with a half-life to reach to the PSS_460_ of 2.6 s after illumination at 460 nm **(Table 1)**. The thermal relaxation rate of the *cis* isomers to *trans* configuration in the dark is especially slow, probably in the order of days, but solubility issues preclude their pre*cis*e determination. In addition, azoglurax photoisomerization is stable and reversible along the multiple UV/blue illumination cycles **(Figure 2B)**.

### 4.3. Azoglurax is a submicromolar, *trans*-active photoswitchable ago-PAM of mGlu_5_

Next, we assessed the pharmacological profile of azoglurax in HEK293 cells expressing the mGlu_5_ receptor using an IP1 accumulation assay.

We evaluated the ability of different concentrations of azoglurax to enhance the dose-dependent activation of mGlu_5_ induced by the agonist quisqualate, in the dark and upon illumination with 365nm light **(Figure 3A and 3B)**. The profile obtained suggests that, similarly to its parent molecule the agonist PAM VU0424465, azoglurax possesses a strong intrinsic agonist activity, as can be seen by the production of IP1 measured in absence of quisqualate. Indeed, the analysis with the Hill equation reveals an increase of the Bottom values, a geometric indication of allosteric agonist activity. However, we did not observe a significant potentiation of quisqualate potency **(Table 2)**.

**Figure 3.**
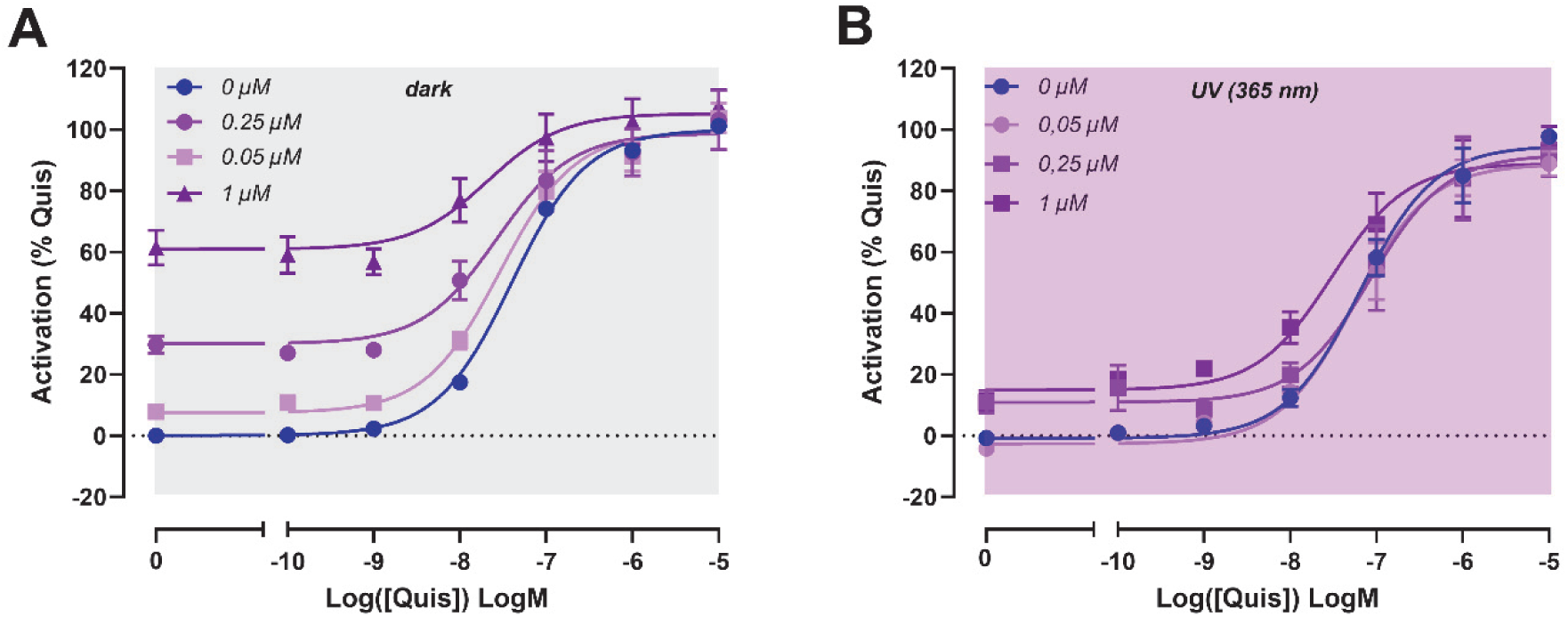
Azoglurax enhancement of mGlu_5_ signaling is decreased upon UV illumination. Receptor activation was determined by measurement of the production of the second messenger inositol phosphate in HEK293 cells *trans*iently expressing mGlu_5_. Dose-dependent potentiation of the response of mGlu_5_ induced by various doses of agonist (quisqualate) by different doses of azoglurax in the dark (**A**) and under UV illumination (**B**). Data are normalized to the maximal efficacy induced by quisqualate. All data are presented as mean ± SEM of at least n=3 experiments (see Table 3 for pEC50, Emax and Ebasal values and number of replicates).

**Table 2.** Summary of potency (pEC50), basal efficacy (Ebasal) and maximal efficacy (Emax) values of quisqualate on mGlu_5_ in presence of various concentrations of azoglurax, in the dark or under UV light. Data are presented as mean ± SEM from n independent experiments (n between brackets) of allosteric modulation of quisqualate-mediated responses by azoglurax, corresponding to the curves displayed in Figure 3. Data are normalized to the maximal efficacy of quisqualate alone.

|  | Dark |  |  | UV Light |  |  |
| --- | --- | --- | --- | --- | --- | --- |
|  | pEC<sub>50</sub> | E<sub>basal</sub> | E<sub>max</sub> | pEC<sub>50</sub> | E<sub>basal</sub> | E<sub>max</sub> |
| Quisqualate | 7.40 ± 0.03 (7) | 0 (7) | 100 (7) | 7.31 ± 0.07 (5) | 0 (5) | 100 (5) |
| + 0.05 μM Azoglurax | 7.56 ± 0.04 (3) | 7.43 ± 0.42 (3) | 99.24 ± 5.87 (3) | 7.24 ± 0.14 (3) | -2.72 ± 0.63 (3) | 89.24 ± 3.97 (3) |
| +0.25 μM Azoglurax | 7.58 ± 0.09 (4) | 29.99 ± 1.26 (4) | 99.09 ± 8.50 (4) | 7.44 ± 0.03 (3) | 10.55 ± 3.59 (3) | 88.42 ± 9.12 (3) |
| +1 μM Azoglurax | 7.55 ± 0.07 (3) | 67.20 ± 5.19 (3) | 109.14 ± 7.18 (3) | 7.46 ± 0.19 (3) | 14.79 ± 1.48 (3) | 96.09 ± 15.85 (3) |

**Table 3.**
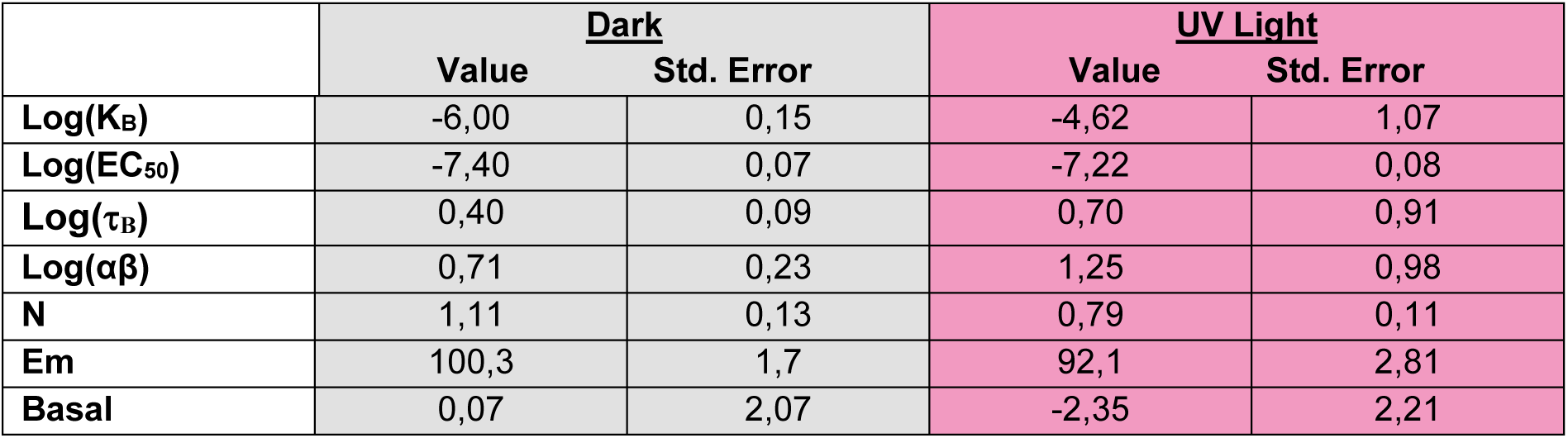
Analysis of the allosteric modulation of quisqualate-mediated responses by azoglurax using the operational model of allosterism. Data obtained from dose-response curves of quisqualate in presence of various concentrations of azoglurax in the dark and under illumination, analyzed with the operational model of allosterism (Equation 1) [51]. Parameter estimates and standard errors produced by global fitting.

The effect of UV light is mainly observed on the agonist activity of azoglurax. For example, upon illumination with 365nm, the production of IP1 induced by 1 μM of azoglurax in absence of quisqualate is decreased from 60% in the dark to 10%. Interestingly, this would be compatible with the activity carried out by the remaining *trans* isomers in solution in these light conditions. The data analysis with the operational model **(table 3)** suggests that the main reason for the diminished activity of *cis* azoglurax could be a reduction of the binding affinity.

### 4.4. Azoglurax is an allosteric agonist that activates mGlu_5_ in absence of agonist

Next, we evaluated the ability of increasing concentrations of azoglurax to enhance the activity of mGlu_5_ induced by a low concentration of the agonist quisqualate (10 nM ≈ EC_20_), in the dark and upon illumination with 365-nm light, using the parent molecule VU0424465 as a control **(Figure 4A)**. In the dark, azoglurax displays a robust and dose-dependent potentiation of agonist-induced activity of mGlu_5_, with a pEC_50_ of 6.56 ± 0.13 (n=11). Upon UV illumination, azoglurax displays a 5-fold shift in potency while its efficacy level is maintained **(Table 4)**. Of note, the potency and efficacy of VU0424465 are not significantly affected by UV illumination.

**Figure 4.**
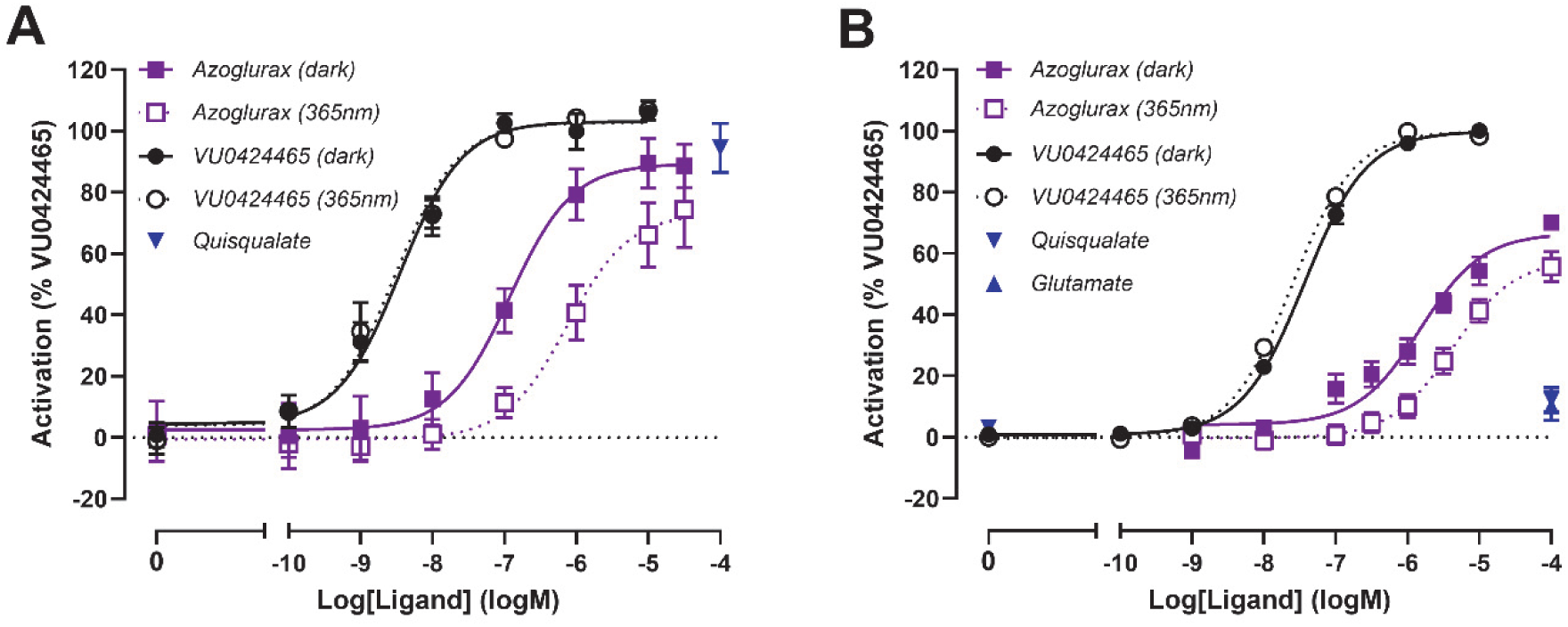
Dose-dependent potentiation of the response of mGlu_5_ and intrinsic agonist activity of azoglurax in absence of agonist. Receptor activation was determined by measurement of the production of the second messenger inositol phosphate in HEK293 cells transiently expressing wild-type mGlu_5_ or an agonist-insensitive mGlu_5_ mutant (mGlu_5_ YADA). Dose-dependent potentiation of the response of mGlu_5_ induced by a single dose of agonist (quisqualate, 10 nM) by different doses of azoglurax and VU0424465, in the dark or under UV illumination (365 nm, 6 mW) on mGlu_5_ wt (**A**) and mGlu_5_ YADA (**B**). Glutamate and quisqualate at high dose (100 µM) are only inducing a very moderate effect on mGlu_5_ YADA. Data are normalized to the maximal efficacy induced by VU0424465. Data are presented as mean ± SEM of at least 3 separate experiments for azoglurax, glutamate and quisqualate and VU0424465 (see Table 4 for potency and efficacy values and number of replicates).

**Table 4.**
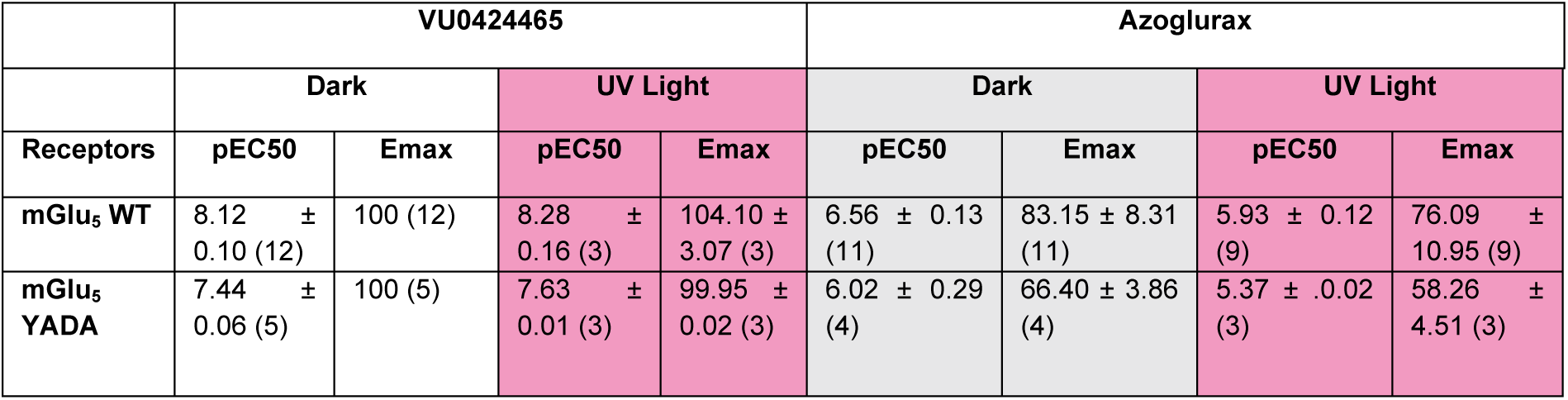
Summary of potency and maximal efficacy of azoglurax and VU0424465 on mGlu_5_ WT and mGlu_5_ YADA, in the dark or under UV illumination. pEC50 and Emax are calculated from the n individual experiments corresponding to the curves displayed in Figure 4. In mGlu_5_ WT, these parameters are determined from the dose-dependent potentiation by azoglurax and VU0424465 of the response induced by 10 nM quisqualate. In mGlu_5_ YADA, they are determined from the dose-dependent activation of this agonist-insensitive mutant by azoglurax and VU0424465 alone. Emax are normalized to the maximal efficacy of VU0424465 measured in the dark and expressed in %. Data are presented as mean ± SEM from n independent experiments (n between brackets).

We cannot completely rule out the possibility that, at least in part, the agonist activity of azoglurax on mGlu_5_ observed in absence of added agonist could result from a potentiation of ambient glutamate in the cell medium. However, it is unlikely that it causes all of the agonist activity because our experimental conditions are designed to reduce the presence of ambient glutamate as much as possible, through the use of GlutaMAX supplement and the co-expression of the high-affinity glutamate transporter EAAC1, as previously described [55].

To further evaluate the intrinsic agonist activity of azoglurax, we thus performed a series of experiments using an mGlu_5_ mutant insensitive to glutamate **(Figure 4B)**. This mutant receptor (mGlu_5_ YADA) possesses two mutations in the agonist-binding site (Y236A and D318A), which greatly impair glutamate and quisqualate action. However, both VU0424465 and azoglurax conserves their ability to activate this receptor, although with a decreased potency and efficacy when compared to that measured on the WT receptor **(Table 5)**. These data confirmed that azoglurax is an allosteric agonist of mGlu_5_.

**Table 5.**
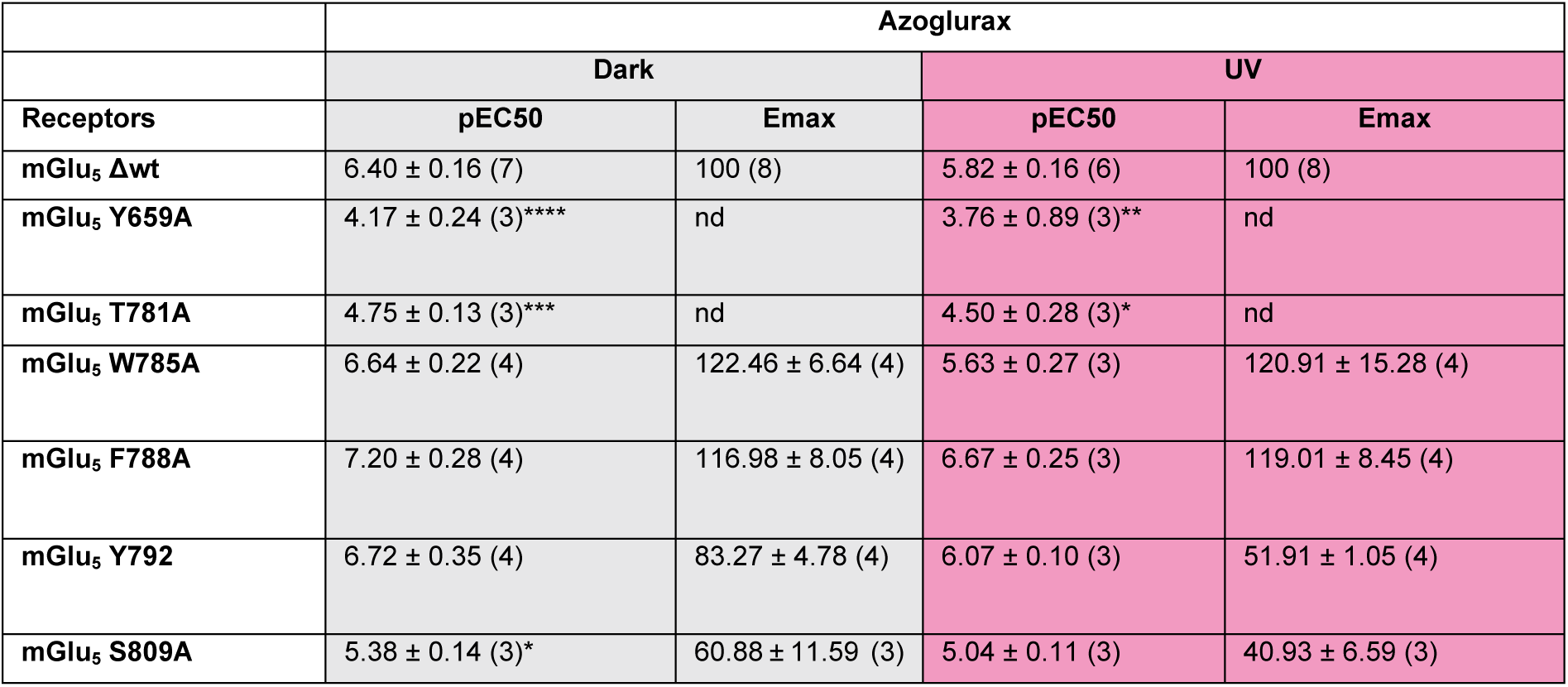
Potency and efficacy of azoglurax on mGlu_5_ wt and binding site mutants. pEC50 ± SEM are calculated from n individual experiments (n between brackets) corresponding to the figure 5, except for the mutant mGlu5 Y659A for which the pEC50 values are extrapolated from dose-response curves. Maximal efficacy values (Emax) are normalized to the Emax of the ligand measured on the mGlu_5_ ΔWT receptor and expressed in %. Statistical analysis: One-way ANOVA comparing pEC50 mutant vs pEC50 ΔWT, followed by a Dunnett post-hoc test. P values: *p<0.05 are shown as *; **p<0.01; ***p<0.001; ****p<0.0001.

| Receptors | Azoglurax |  |  |  |
| --- | --- | --- | --- | --- |
|  | Dark |  | UV |  |
|  | pEC<sub>50</sub> | E<sub>max</sub> | pEC<sub>50</sub> | E<sub>max</sub> |
| mGlu<sub>5</sub> Δwt | 6.40 ± 0.16 (7) | 100 (8) | 5.82 ± 0.16 (6) | 100 (8) |
| mGlu<sub>5</sub> Y659A | 4.17 ± 0.24 (3)**** | nd | 3.76 ± 0.89 (3)** | nd |
| mGlu<sub>5</sub> T781A | 4.75 ± 0.13 (3)*** | nd | 4.50 ± 0.28 (3)* | nd |
| mGlu<sub>5</sub> W785A | 6.64 ± 0.22 (4) | 122.46 ± 6.64 (4) | 5.63 ± 0.27 (3) | 120.91 ± 15.28 (4) |
| mGlu<sub>5</sub> F788A | 7.20 ± 0.28 (4) | 116.98 ± 8.05 (4) | 6.67 ± 0.25 (3) | 119.01 ± 8.45 (4) |
| mGlu<sub>5</sub> Y792 | 6.72 ± 0.35 (4) | 83.27 ± 4.78 (4) | 6.07 ± 0.10 (3) | 51.91 ± 1.05 (4) |
| mGlu<sub>5</sub> S809A | 5.38 ± 0.14 (3)* | 60.88 ± 11.59 (3) | 5.04 ± 0.11 (3) | 40.93 ± 6.59 (3) |

### 4.5. Azoglurax shares a common binding mode with its parent molecule VU0424465

We investigated the molecular determinants of azoglurax binding to mGlu_5_ using molecular modelling coupled to site-directed mutagenesis. To that aim, *trans-*azoglurax was docked to the allosteric pocket of the recently reported high-resolution cryoEM structure of mGlu_5_ bound to VU0424465 [12], which resulted in a similar binding pose of *trans-*azoglurax and VU0424465 **(Figure 5A)**. Then, we performed molecular dynamics (MD) simulations of *trans*-azoglurax, *cis*-azoglurax and VU0424465 in the allosteric pocket of mGlu_5_ 7TM **(movies)**. *Cis*-azoglurax was obtained by switching the -N=N- double bond of *trans*-azoglurax in the docked model. *Trans*-azoglurax maintained a nearly identical binding pose to VU0424465 during the simulations **(Figure 5A** and **Movie 1)**. However, it showed larger fluctuations, consistent with its lower potency observed *in vitro*. On the other hand, *cis*-azoglurax was highly mobile during the same simulation course (**Movie 1**), indicating low binding affinity.

**Figure 5:**
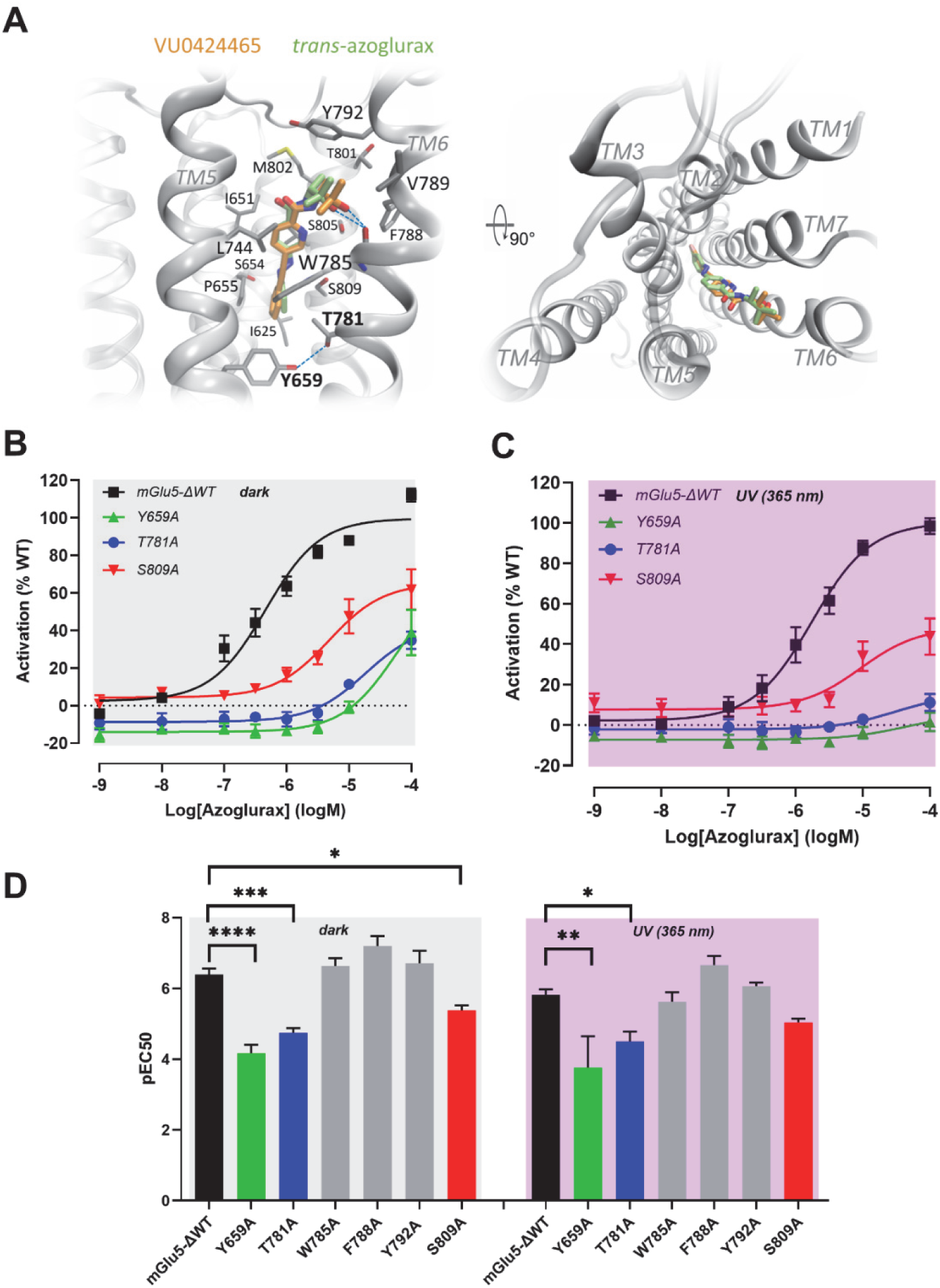
Binding mode of azoglurax. **A.** Docking and MD simulations of *trans-*azoglurax vs VU0424465 in the cryoEM structure of mGlu_5_ bound to VU0424465 from Cannone et al. [12] show that the two ligands share a similar binding pose. VU0424465 exhibited a more stable hydrogen bond (blue dashed lines) with W785 backbone. **B, C.** Dose-response curves of azoglurax on mGlu_5_ ΔWT, mGlu_5_ T781A, mGlu_5_ F788A and mGlu_5_ S809A in the dark (B) and under light (365 nm, 8 mW)(C). **D.** Mean pEC50 values of azoglurax on the different mutants of mGlu_5_ binding pocket. Similar residues are key for azoglurax and VU0424465 pharmacological activity. Receptor activation was determined by measurement of the production of the second messenger inositol phosphate in HEK293 cells transiently expressing mGlu_5_ ΔWT and mutants. Data are mean ± SEM of at least n=3 experiments (see Table 4 for pEC50 values and number of replicate).

In order to probe the docking of azoglurax, we then used site-directed mutagenesis to substitute the amino acids surrounding azoglurax by alanine and evaluate its ability to enhance 10 nM quisqualate activity on these mutants. Azoglurax activity was assessed in the dark or under UV illumination. Functional data are summarized in **Table 5**. The dose-response curves of azoglurax on the mutants that caused significant functional consequences are displayed in **Figure 5B-C**. The activity of azoglurax was significantly altered in three out of the six mutants: Y659A, T781A and S809A for which its potency and efficacy are reduced with regard to the wild type receptor, both in the dark and under UV illumination. Y659, T781 and S809 outline the narrow bottom of the allosteric pocket, where Y659 and T781 form a hydrogen bond (**Figure 5A**). These three residues likely maintain the pocket shape and depth. Thus, mutating them to alanine had significant impact on both *trans*- and *cis*-azoglurax. Interestingly, as previously shown and corroborated herein [12, 56], these amino acids are also inducing a significant reduction of the activity of VU0424465 **(Table 5)**, further confirming that azoglurax and its parent molecules are sharing a similar binding mode. The other three mutations, W785A, F788A and Y792A, showed insignificant impact on the potency or efficacy of azoglurax. These residues are located in the upper half of the allosteric pocket on TM6. They appeared more dynamic during the MD simulations due to the TM6 movements. This may explain their lack of impact upon mutation despite the close proximity to the ligand.

## 5. Discussion

The mGlu_5_ receptor is widely expressed in the brain where it plays an important role in synaptic plasticity. It is also a therapeutic target of interest in various nervous system pathologies, such as anxiety, chronic pain, neurodegenerative diseases and mental disorders. This receptor has been the target of numerous drug discovery programs, leading to the development of many selective allosteric ligands. However, despite great promise, there are no drugs on the market targeting mGlu_5_ to date, underlining the importance of continuing investigations into this target. Photopharmacology, through the fine control of drug action in time and space by light, is potentially a new route to targeted, localized and effective treatments with limited off-target side effects [25]. Upstream, photopharmacology also offers the possibility of studying and understanding the function of a biological target with great spatiotemporal precision. Although various photopharmacology tools have been developed for this receptor over the last few years, they are all NAMs and to date there has been no photocontrolable activator for mGlu_5_ [57]. Here we present the first freely-diffusible photoswitchable ago-PAM of mGlu_5_, named azoglurax, which enables the optical control of receptor activation.

We designed and synthesized azoglurax through an azologization strategy [27] of the well-known mGlu_5_ ago-PAM VU0424465 [36]. This strategy was chosen instead of azo extension because it has proven efficient in the past for generating photoswitchable allosteric modulators for mGluRs, such as Alloswitch-1 [31], Optogluram [58] and their derivatives [32, 59, 60]. As expected, the azo incorporation conferred to azoglurax the ability to reversibly photoisomerize upon illumination at appropriate wavelengths. The optimal wavelength for *trans* to *cis* photoisomerization is in the UV range (365 nm), while the reverse *cis* to *trans* photoswitch is in the blue range (460 nm). The *trans* to *cis* photoconversion following illumination at 365 nm is quite limited, with a solution at the photostationary state containing 70% of *cis*-isomer, while following illumination at 460 nm the photostationary state contains 84% of the *trans* isomer.

On the pharmacological point of view, the azologization process affected both the potency and efficacy of the ligands, as evidenced by the ≈50-fold shift in potency and the ≈25% decrease in efficacy compared to VU0424465. However, since VU0424465 is an exceptionally potent mGlu_5_ agonist PAM, azoglurax retains a submicromolar potency. Interestingly, azoglurax conserved the allosteric agonist activity of VU0424465 as well as its binding mode. Both ligands display a strong intrinsic agonist activity, inducing mGlu_5_ signaling in absence of agonist and direct activation of an agonist-insensitive mGlu_5_ mutant. In the present study, molecular modeling, docking and mutagenesis reveal that both ligands bind in the same binding pocket within the 7TM of the receptor. Notably, the same two key residues located at the bottom of the pocket, Y659 in the TM3 and T781 in the TM6 are important for VU0424465 and azoglurax potencies [12, 56, 61]. Altogether, these data suggest that the ago-PAM activity of both azoglurax and VU0424465 probably occur through the same mechanisms. However, the azobenzene moiety of azoglurax is very similar to that of alloswitch-1 and they may interact with similar residues as reported for alloswitch-1 in the inactive state of the receptor [9]. Stabilizing the same set of residues Y659, S809, T781, W785) may in part contribute to the lower potency of azoglurax in its *trans* conformation [9, 12].

From a photopharmacological point of view, ago-PAM activity of azoglurax is higher in the *trans* state (in absence of light) than under illumination with 365-nm light to obtain a mixture enriched with the *cis* configuration, as most of the azobenzene-containing photoswitchable ligands described so far and in particular those acting on mGluRs [57]. The photoisomerization of azoglurax can likely occur both in solution and while bound in the pocket, similarly to what is observed for the photoswitchable mGlu_5_ NAMs Alloswitch-1 and MCS-331 [59, 62]. When occurring in the receptor, the *trans* to *cis* photoisomerization destabilizes the binding of azoglurax to the receptor as shown by MD simulation, which results in the decrease of its affinity for the receptor. A theoretical pEC50 if only *trans*- azoglurax was active can be extrapolated based on the 70/30 *cis*/*trans* ratio determined under illumination at 365 nm. This theoretical pEC50 is very close to the pEC50 determined experimentally (extrapolated pEC50: 6.44; experimental pEC50: 6.56), suggesting that the remaining *trans*- azoglurax is mostly responsible for the activity observed and that *cis*-azoglurax has a very low potency or fully inactive. The development of derivatives with improved photoswitching properties may be of interest in the future. To improve the *trans* to *cis* photoconversion and the photostationary state of azoglurax derivatives, several solutions are existing such as the addition of substituents to the aromatic rings to induce an effective separation of bands or their replacement by heteroaryls, such as pyriazole, to induce a push-pull effect, which could also help to improve a photostationary state richer in *cis*-isomer [64]. Besides, the addition of tetra-*ortho*-substituents to the azobenzene can allow a shift of the isomerisation wavelengths to more penetrant and less harmful wavelengths in the visible range [63].

In conclusion, with the photoswitchable agonist PAM azoglurax presented here, we add a novel tool to the photopharmacological toolbox of mGlu_5_, enabling the optical control of receptor activation. Azoglurax could be useful for studying the mechanisms of action of mGlu_5_ *in vitro* or *in cellulo* with good spatiotemporal resolution. For example, this photoswitchable activator may be of interest to explore the signaling of mGlu_5_ in native cells such as neurons and astrocytes which naturally express this receptor or to assess the structural dynamics and activation mechanisms of mGlu_5_ *in vitro*.

## 6. Acknowledgements

The authors thank Lourdes Muñoz from SIMChem (IQAC-CSIC) for the use of analytical support and helpful discussions, Charleine Zussy and Mathieu Vitalis for helpful discussions, and the staff of Montpellier Biocampus Arpege facility for their technical help. MD simulations were performed using supercomputing resources from GENCI-TGCC France. (grant A0150714630 2023-2024 to XC). This work was funded by the French National Research Agency (ANR) under the “Investissements d’avenir” program I-Site MUSE with the reference ANR-16-IDEX-0006 to AD, LG and CG, the reference ANR-17-NEU3-0001 under the frame of Neuron Cofund to RBT, AL and CG, and the reference ANR-20-CE11-019 to GL. This work was also financially supported by the Fondation pour la Recherche Médicale, grant number «MND202003011477», to IT, LG and CG and by the International Emerging Action (IEA) from the Centre National de la Recherche Scientifique (CNRS) to CG and AL, Ministerio de Ciencia e Innovación, Agencia Estatal de Investigación, and ERDF - A way of making Europe (PID2020-120499RB-I00, and PIE-RYC2020--029485-I) to A.L. and X.R., by the Agència de Gestió d’Ajuts Universitaris i de Recerca (2021 SGR 00508) and Tecnio association (URD316) to A.L. and by the Spanish National Research Council (20228AT014) to X.R. and European Union’s Horizon 2020 research and innovation programme under Marie Skłodowska-Curie grant agreement no. 801342 (Tecniospring INDUSTRY, TECSPR19-1-0062) and the Government of Catalonia’s Agency for Business Competitiveness (ACCIÓ) to XG-S.

## 7. Authors contributions CREDIT

Conceptualization, GL, AL and CG; Methodology, XGS, FM, XC, GL, AL, and CG; Investigation, AD, RBT, FM, XC, IT, XR, FM, and CG; Resources, AL, XC, GL and CG; Writing original draft: AD and CG; Writing review and editing: all authors; Visualization, AD and CG; Funding Acquisition: CG, LG, GL, and AL; Supervision, XGS, GL, AL and CG

## Abbreviations

mGluRs: metabotropic glutamate receptors
PAM: positive allosteric modulator
NAM: negative allosteric modulator
ago-PAM: PAM agonist
7TM: seven transmembrane alpha-helical domain
IP_1_: inositol monophosphate
(PSS): photostationary state

## Supplemental materials

Videos file: Movie REST2 MD simulations of ligands movement in allosteric pocket.mp4

Legend : Movements of each ligand in the allosteric pocket during 50 ns × 64 replicas (3.2 µs) of REST2 MD simulations.

